# The hepatokine FGL1 regulates hepcidin and iron metabolism during the recovery from hemorrhage-induced anemia in mice

**DOI:** 10.1101/2023.04.06.535920

**Authors:** Ugo Sardo, Prunelle Perrier, Kevin Cormier, Manon Sotin, Aurore Desquesnes, Lisa Cannizzo, Marc Ruiz-Martinez, Julie Thevenin, Benjamin Billoré, Grace Jung, Elise Abboud, Carole Peyssonnaux, Elizabeta Nemeth, Yelena Z. Ginzburg, Tomas Ganz, Léon Kautz

**Affiliations:** IRSD, Université de Toulouse, INSERM, INRAE, ENVT, Univ Toulouse III - Paul Sabatier (UPS), Toulouse, France; Icahn School of Medicine at Mount Sinai, New York, NY, USA; Departments of Medicine, David Geffen School of Medicine at UCLA, Los Angeles, CA; Departments of Pathology, David Geffen School of Medicine at UCLA, Los Angeles, CA; Institut Cochin, INSERM, Centre National de la Recherche Scientifique (CNRS), Université de Paris, Paris, France; Laboratory of Excellence GR-Ex, Paris, France

**Author notes:** US and PP contributed equally.

## Abstract

As a functional component of erythrocyte hemoglobin, iron is essential for oxygen delivery to all tissues in the body. The liver-derived peptide hepcidin is the master regulator of iron homeostasis. During anemia, the erythroid hormone erythroferrone regulates hepcidin synthesis to ensure adequate supply of iron to the bone marrow for red blood cells production. However, mounting evidence suggested that another factor may exert a similar function. We identified the hepatokine FGL1 as a previously undescribed suppressor of hepcidin that is induced in the liver in response to hypoxia during the recovery from anemia and in thalassemic mice. We demonstrated that FGL1 is a potent suppressor of hepcidin *in vitro* and *in vivo*. Deletion of *Fgl1* in mice results in a blunted repression of hepcidin after bleeding. FGL1 exerts its activity by direct binding to BMP6, thereby inhibiting the canonical BMP-SMAD signaling cascade that controls hepcidin transcription.

**Key points:** **1/ FGL1 regulates iron metabolism during the recovery from anemia.**

**2/ FGL1 is an antagonist of the BMP/SMAD signaling pathway.**

## INTRODUCTION

Anemia, defined as a decreased number of functional red blood cells, is a major cause of morbidity and mortality affecting one-third of the worldwide population^1^. Many conditions including iron deficiency, bleeding, infections and genetic disorders may cause anemia. Iron is an essential functional component of erythrocyte hemoglobin, and sustained delivery of iron for erythropoiesis in the bone marrow is required to maintain the erythron and tissue oxygenation^2, 3^. Iron is released from iron recycling macrophages, enterocyte and hepatocytes by the sole iron exporter ferroportin. The liver-derived hormone hepcidin regulates body iron content by binding to ferroportin, occluding it and triggering its degradation^4, 5^.

Hepcidin synthesis is predominantly regulated by the canonical BMP-SMAD signaling pathway and the bone morphogenetic proteins BMP 2 and BMP 6^6–8^. Binding of BMP 2/6 to a large receptor complex leads to the phosphorylation of effectors SMAD 1, 5 and 8 that translocate into the nucleus to activate hepcidin transcription^9^. Hepcidin expression is rapidly suppressed by the erythroid regulator erythroferrone (ERFE) in conditions associated with expanded erythropoiesis such as anemia caused by bleeding or inflammation^10, 11^. Conversely, excessive release of ERFE in inherited anemias (e.g. beta thalassemia, congenital dyserythropoietic anemia) or in myelodysplastic syndromes^12–14^ causes iron overload and, if untreated, lethal clinical complications. In response to erythropoietin (EPO), ERFE is secreted by erythroid precursors in the bone marrow and the spleen, and sequesters BMP 2/6 in the liver perisinusoidal space to prevent BMP receptor engagement and inhibit the signaling cascade directing hepcidin expression^15^.

Although ERFE is essential for the suppression of hepcidin within the first hours following an erythropoietic stress, *Erfe-*deficient mice recover from anemia induced by hemorrhage and chronic inflammation^10, 11^. In thalassemic mice, ablation or neutralization of ERFE^16, 17^ raises hepcidin levels and mitigates the systemic iron overload. However, restoration of physiological levels of hepcidin is not sufficient to correct the iron overload and hepcidin synthesis remains inappropriately low relative to the liver siderosis. Collectively, these data indicated that one or more ERFE-independent mechanisms repressed hepcidin during anemia.

We therefore examined hepcidin regulation during the recovery from hemorrhage-induced anemia in WT and *Erfe*-deficient mice and confirmed the ERFE-independent repression of hepcidin during anemia. Here we describe the identification of a new BMP antagonist and hepcidin suppressor: the liver produced hepatokine Fibrinogen-like 1 (FGL1).

## METHODS

### Animal models

*Erfe*-/- and WT controls on a C57BL/6J background were bred and housed in a specific and opportunistic pathogen-free barrier facility in the animal facilities of INSERM US006. *Fgl1-/-* and WT controls on a C57Bl/6N background were obtained from The European Mouse Mutant Archive (EMMA), bred by Janvier labs (Le Genest St Isle) and transferred in the animal facilities of INSERM US006 at the age of 4-5 weeks. Mice were housed under a standard 12- hour light/dark cycle with water and standard laboratory mouse chow diet (Ssniff, 200 mg iron/kg) *ad libitum*, in accordance with the European Union guidelines. The study was approved by the Midi-Pyrénées Animal Ethics Committee. Thalassemic mice and mice treated with Vadadustat were provided by Yelena Ginzburg^18^ and Tomas Ganz^19^ respectively. Expression data from *Vhlh^lox/lox^/AlbuminCre* mice were provided by Carole Peyssonnaux^20^. To study the recovery from anemia, mice were phlebotomized by a retro-orbital puncture (500µL) and analyzed after 1 to 6 days. Disruption of the erythroid compartment was achieved by exposing mice to a sublethal dose of X-ray (400 rads) and mice were phlebotomized 48 hours later. Surgical ablation of the spleen was performed on 7–8-week-old WT and *Erfe-/-* mice. Mice were allowed to rest for 7 days before phlebotomy. A subset of WT mice was given a single dose of EPO (200U) and were analyzed 12, 15, 18 or 20 hours later. Seven-week-old C57Bl/6J mice fed for two weeks with an iron adequate diet (Ssniff, 50 mg/kg) were injected intra-peritoneally with recombinant FGL1, Fc fragment (hIgG2) or saline at a dose of 10mg/kg and analyzed after 6 hours. For all mice, tissues were harvested and divided into flash frozen sample in liquid nitrogen for RNA, protein and iron measurements and in 4% formalin for paraffin embedding. Male mice were preferentially studied unless otherwise specified.

### Production of recombinant FGL1

Mouse FGL1 cDNA sequences (full length, N-terminal domain, globular domain) were cloned into pFUSEN-hG2Fc plasmid (InvivoGen) with the following modifications: vector signal sequence (from Interleukin-2) was used instead of the native, followed by the Fc fragment of human IgG2. Recombinant proteins were produced in suspension culture in Freestyle 293F cells (Life Technologies) transiently transfected using FectroPro reagent (Polyplus). Supernatants from cells overexpressing Fc-tagged FGL1 proteins were collected after 5 days and supplemented with protease inhibitor cocktail (Sigma). Recombinant proteins were purified using Hitrap Protein A HP column on an ÄKTA pure chromatography system (GE healthcare) and eluted with 0.1M Glycine pH 3.5. The eluted fractions were concentrated using centrifugal concentrators Spin-X UF 20 (Corning), and recombinant FGL1 proteins were suspended in a saline solution (0.9% NaCl). Protein purity and concentration were determined using Coomassie Imperial Protein Stain and Pierce bicinchoninic acid protein assay (Thermo Fisher Scientific).

### Mouse ERFE Immunoassay

Human recombinant monoclonal antibodies to mouse ERFE were produced by Bio-rad using the HuCAL technology. High binding 96 well plate (Corning) was coated overnight at 4°C with 100 µL/well of 2 µg/ml capture antibody diluted in 50 mM sodium carbonate buffer pH 9.6. Plate was washed (TBS, 0.05% Tween 20) and blocked for an hour with 300 µL/well blocking buffer (PBS, 0.2% Na casein, 0.05% Tween 20, 0.1 M NaCl) at room temperature. Recombinant mouse ERFE standard was serially diluted to 10, 5, 2.5, 1.25 and 0. 625 ng/ml. Serum samples diluted in PBS and standards diluted in PBS + 5% BSA were incubated for 1 hour incubation at room temperature. Plate was washed and incubated for 1 hour with 100µL/well of biotinylated detection antibody at 0.5µg/ml in PBS + 5% BSA. Plate was washed, and incubated for 45 minutes with 100µl/well of 1/5000 Neutravidin-HRP (Pierce) in PBS + 5% BSA. Plate was developed with 100 µL/well Supersensitive TMB substrate (Thermofisher) in the dark at room temperature, the reaction was stopped by adding 50µL of 2N sulfuric acid, and the absorbance was measured at 450 nm.

### Measurement of iron and hematological parameters

Serum iron concentration was determined by iron direct method (ferene, Biolabo, 92108) and transferrin saturation was deduced by measuring the unsaturated iron binding capacity (UIBC, Biolabo, 97408). Liver iron content was determined as previously described^21^. Complete blood count was performed with a Cell-Dyn Emerald hematology analyzer (Abbott).

### Western Blot Analysis

Liver proteins were extracted by physical dissociation (Ultra-turrax, IKA) in PEB Buffer (150 mM NaCl, 50 mM Tris-HCl, 5mM EDTA, 1% NP-40) containing proteases (cOmplete™, Roche) and phosphatase (Phosphatase Inhibitor Cocktail 2, Sigma) inhibitors. Hep3B cells were lyzed in RIPA Buffer (Thermo Fischer Scientific) containing protease and phosphatase inhibitors. Freshly extracted proteins were diluted in Laemmli buffer 2x (Sigma), incubated 10 min at 95°C, subjected to SDS-PAGE and electroblotted to nitrocellulose membrane (Bio-rad). Membranes were blocked 1h with 5% of non-Fat dry milk (NFDM, Cell signaling) diluted in TBS-T buffer (10mM Tris-HCl, pH 7.5, 150mM NaCl, 0.15% Tween 20) and incubated overnight at 4°C with phospho-Smad5 (Ser463/465, Abcam, ab92698, 1/2000) or 2 hours at RT with antibody to Smad5 (Abcam, ab40771, 1/5000) diluted in TBS-T buffer 5% BSA. Loading was determined using antibodies to GAPDH (Cell signaling, D16H11, 1/10 000) or Vinculin (Cell signaling, 4650, 1/20 000) diluted in TBST- NFDM (5%) (2h, RT). Incubation with primary antibody was followed by 3 washes and membranes were incubated 2 hours with goat anti-Human IgG (Novus biological, NBP1-75006, 1/10000), goat anti-rabbit IgG (Cell signaling, 7074, 1/10000) or horse anti-mouse IgG (Cell signaling, 7076, 1/10000) secondary antibodies conjugated with HRP and diluted in TBST- NFDM (5%). Enzyme activity was developed using ECL prime reagent (GE Healthcare) on ChemiDoc XRS+ imaging system.

### Pull down assay

One microgram of Fc tagged recombinant proteins (Fc alone, FGL1 full length, FGL1 globular and FGL1 Nter) was incubated overnight at 4°C with protein A magnetic beads (Dynabeads, Pierce) in NETN buffer (20 mM Tris-HCl pH 8.0, 0.5% NP-40, 100 mM NaCl, 1 mM EDTA pH 8.0, Protease inhibitor cocktail) with or without 500 ng of hBMP6 (Biotechne). Proteins were eluted using Laemmli buffer and analyzed by western blot using Goat anti-Human IgG Fc fragment Secondary Antibody [HRP] (Novus biological NBP1-75006) or anti-BMP6 antibody (R&D systems, AF6325).

### Cell treatment

Hep3B and HepG2 cells were culture in Dulbecco’s modified Eagle medium-high glucose GlutaMAX, 10% fetal bovine serum, 1% penicillin-streptomycin. Cells were plated 24 hours before treatments and treated for 6 hours. Hepatocytes were isolated from wild-type C57BL/6J mice by a portal vein collagenase perfusion method as previously described^22^. Cells were incubated overnight (15 hours) in fresh Williams E Medium (Gibco) supplemented with 200 µM L-glutamine, 10% FBS. Hep3B and HepG2 cells and primary hepartocytes were treated with 10 or 25 ng/ml of BMP6 (PeproTech) or BMPs 2, 4, 7, (R&D Systems) and with Fc (hIgG2), Fc-FGL1 full length (FL), its N-terminal (Nter) or globular (glob) domains for 6 hours. For hypoxia experiments, cells were maintained in a hypoxia chamber (Whitley, H35 Hypoxystation) with 2% of oxygen for 15 hours or in a conventional CO2 incubator in presence of 1mM DMOG (Dimethyloxalylglycine, N-(Methoxyoxoacetyl)-glycine methyl ester, Sigma).

### Quantification of mRNA levels

Total RNA from mouse tissues was extracted by Trizol (MRC) / Chloroform (Sigma) method. Complementary cDNA was synthetized using M-MLV Reverse transcriptase (Promega). Messenger RNA (mRNA) expression levels were assessed by quantitative polymerase chain reaction (RT-qPCR) using Takyon SYBR green (Eurogentec) (primers indicated in supplemental table 1) and run in duplicate on a LightCycler480 (Roche) apparatus. Transcript abundance was normalized to the reference gene *Hprt* and represented as a difference between reference and target genes within each group of mice (−ΔCt) ± standard error of the mean (SEM). Data from *Vhlh^lox/lox^/AlbuminCre* mice were normalized to the reference gene *36B4*.

### Microarray

Total RNA from mouse liver and bone marrow from *Erfe*-/- mice at t= 0, 24 and 48h hours after phlebotomy was extracted using Trizol (MRC) / Chloroform (Sigma). RNA quality was assessed on RNA 6000 Nano chips using a 2100 Bioanalyzer (Agilent Technologies). Gene-level expression profiling of livers and bone marrows from phlebotomized mice were performed at the GeT-TRiX facility (GénoToul, Génopole Toulouse Midi-Pyrénées) using Agilent Sureprint G3 Mouse GE v2 microarrays (8×60K, design 074809) following the manufacturer’s instructions. For each sample, Cyanine-3 (Cy3) labeled cRNA was prepared from 200 ng of total RNA using the One-Color Quick Amp Labeling kit (Agilent Technologies) according to the manufacturer’s instructions, followed by Agencourt RNAClean XP (Agencourt Bioscience Corporation, Beverly, Massachusetts). Dye incorporation and cRNA yield were checked using Dropsense 96 UV/VIS droplet reader (Trinean, Belgium). 600 ng of Cy3-labelled cRNA were hybridized on the microarray slides following the manufacturer’s instructions. Immediately after washing, the slides were scanned on Agilent G2505C Microarray Scanner using Agilent Scan Control A.8.5.1 software and fluorescence signal extracted using Agilent Feature Extraction software v10.10.1.1 with default parameters.

Microarray data and experimental details are available in NCBI’s Gene Expression Omnibus^23^ and are accessible through GEO Series accession number GSE229041 (https://www.ncbi.nlm.nih.gov/geo/query/acc.cgi?acc=GSE229041). Expression data were analyzed using R (Rv3.1.2)-bioconductor and iDEP (Integrated Differential Expression and Pathway analysis)^24^ by comparing control mice (n=4) to mice 24h (n=4) and 48h (n=4) after phlebotomy.

### Statistical analysis

The statistical significances were assessed by Student’s t-test, one or two-way analysis of variance (ANOVA) using Prism 9 (GraphPad).

## RESULTS

### ERFE-independent repression of hepcidin during anemia

We first delineated the timeline of the recovery from anemia induced by phlebotomy (500µl) in WT and *Erfe*-deficient mice. In both genotypes, we observed that hemoglobin levels decreased for three days after phlebotomy and that mice had recovered normal hemoglobin levels by day 6 (Figure 1A). *Epo* mRNA expression in the kidney was induced on the first two days after phlebotomy in WT and *Erfe-/-* mice and progressively returned to normal after 6 days (Figure 1B). *Erfe* mRNA expression was maximally increased in the bone marrow and spleen after 24 hours and progressively decreased over 6 days. (Figure 1C). Serum ERFE levels were likewise the highest 24h after phlebotomy but decreased below detection levels within 3-4 days (Figure 1D). However, a significant reduction in liver hepcidin mRNA expression was maintained for 1-5 days after phlebotomy in WT mice (Figure 1E). In *Erfe-/- mice*, *Hamp* mRNA expression did not decrease at 24 hours but 2-3 days after phlebotomy dropped to levels comparable to those of WT mice before returning to normal after 5-6 days (Figure 1E). Similar results were observed in phlebotomized female mice (Supplemental Figure 1). Liver *Id1* and *Smad7* mRNA expression was unchanged throughout the time course (Figure 1F-G). We did not detect any statistically significant increase in *Gdf15*^25^ and *Twsg1*^26^ mRNA expression in the marrow and the spleen of phlebotomized WT and *Erfe-/-* mice (Supplemental Figure 2). Serum hepcidin concentration mirrored liver *Hamp* mRNA expression 1 and 2 days after phlebotomy in WT and *Erfe-/-* mice (Supplemental Figure 3A). We therefore decided to focus on the mechanisms triggered within 24-48h after phlebotomy that could contribute to hepcidin suppression in *Erfe-/-* mice. The changes in hepcidin synthesis in *Erfe-/-* mice occurred without any variation in serum iron concentration, transferrin saturation and liver iron content 1 and 2 days after bleeding compared to control mice (Supplemental Figure 3B, C, D). Similarly, SMAD5 phosphorylation was not decreased in the liver of *Erfe-/-* mice 2 days after phlebotomy (Supplemental Figure 3E-F). These data indicate that by 48 hours after phlebotomy hepcidin expression is negatively regulated by an ERFE-independent mechanism.

**Figure 1:**
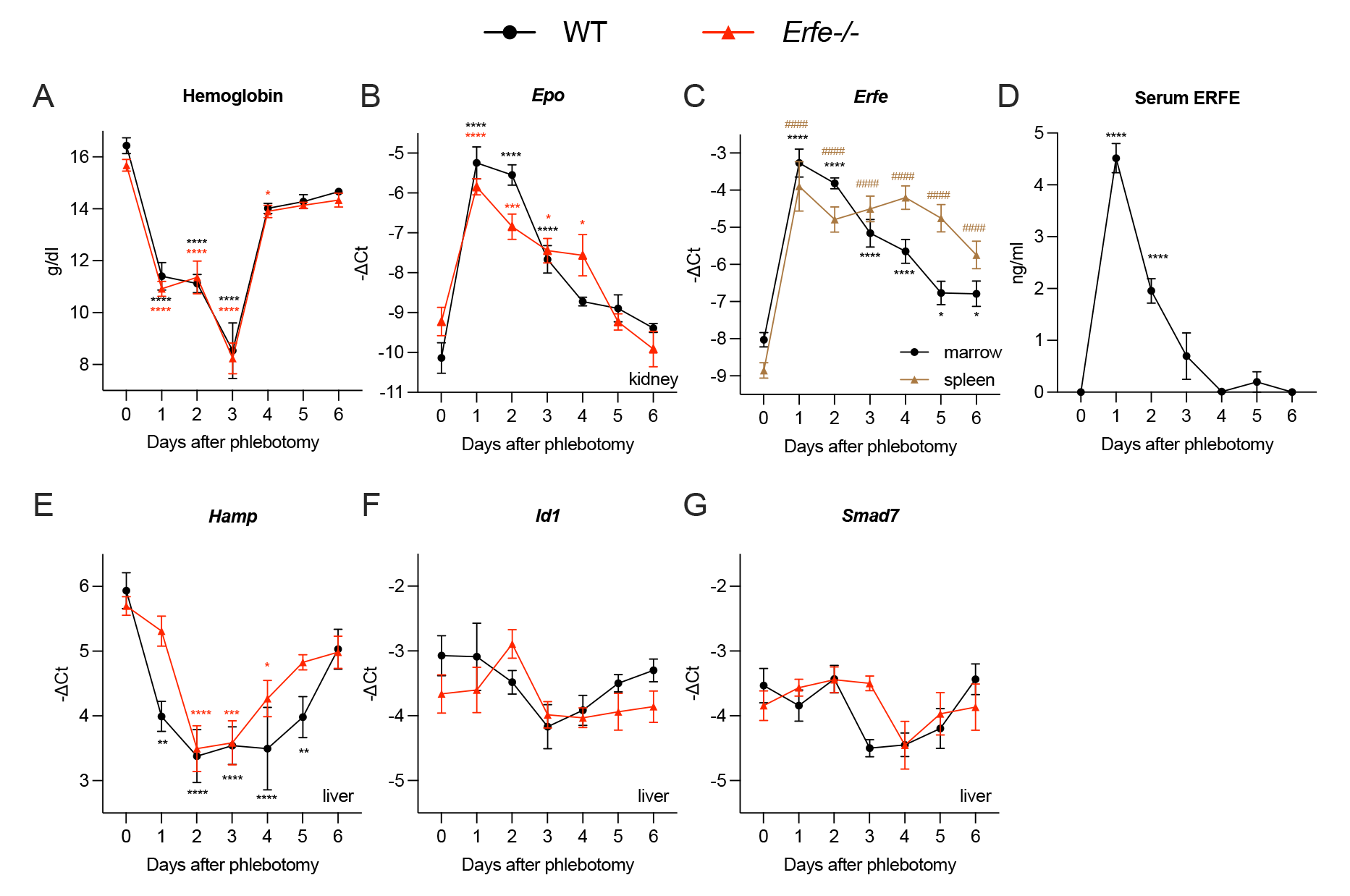
Recovery from hemorrhage-induced anemia in WT and *Erfe-/-* mice. (A) Hemoglobin levels of 7-9 week-old WT (black) and *Erfe-/-* (red) male mice 0, 1, 2, 3, 4, 5 and 6 days after phlebotomy (500µl). mRNA expression of *Epo* in the kidney (B) of phlebotomized WT and *Erfe-/-* mice. Time course of *Erfe* mRNA expression in the bone marrow and spleen (C) and serum ERFE concentration (D) in phlebotomized WT mice. mRNA expression of *Hamp* (E), *Id1* (F) and *Smad7* (G) in the liver of phlebotomized WT and *Erfe-/-* mice. Data shown are means ± s.e.m and were compared for each time point to values for control mice at t = 0 (n = 5 to 8) for each genotype by Two-way ANOVA and corrected for multiple comparisons by Holm-Šidák method. No statistically significant difference was observed between genotypes. \*\*\*\**P* < 0.0001, \*\*\**P* < 0.001, \*\**P* < 0.01, \**P* < 0.05.

### Testing the potential contribution of an erythroid regulator

To determine whether the suppression of hepcidin is mediated by another erythroid regulator^27, 28^, we disrupted the erythroid compartment by irradiation and tested the response of hepcidin to bleeding in WT and *Erfe-/-* mice. Phlebotomized WT and *Erfe-/-* mice not subjected to radiation showed increased reticulocytosis 48h after phlebotomy whereas irradiated mice exhibited a severe decrease in circulating reticulocytes (Figure 2A). Decreased reticulocytes production was accompanied a reduced expression of erythroid markers *Gypa* and *Tfr1* in the bone marrow of irradiated WT and *Erfe-/-* mice compared to their respective controls thus confirming the successful depletion of the erythroid compartment (Supplemental figure 4). Hemoglobin levels were decreased in all phlebotomized groups and to a greater extent in irradiated mice but no difference was observed between WT and *Erfe-/-* mice (Figure 2B). Hepatic hepcidin mRNA expression was repressed in control WT and *Erfe-/-* mice but not in irradiated mice 48 hours after phlebotomy (Figure 2C). However, bone marrow depletion and decreased iron consumption causes an increase in circulating iron concentration which stimulates hepcidin expression and could hinder the effect of competing signals^27^. Interestingly, the recovery from anemia was paralleled by an increase in *Gypa* mRNA expression in the spleen and the liver 1-6 days after phlebotomy compared to control mice (Figure 2D). *Hamp* mRNA expression inversely correlated with *Gypa* mRNA expression in the liver and the spleen (Figure 2E) suggesting that an hepcidin suppressor could be released from these tissues. We thus performed a surgical ablation of the spleen of WT and *Erfe-/-* mice and evaluated the response to bleeding. However, splenectomized WT and *Erfe-/-* mice exhibited reduced liver *Hamp* mRNA expression 48 hours after phlebotomy indicating that the hepcidin suppressive factor did not originate from the spleen (Figure 2F). To explore the liver and bone marrow responses to anemia, we therefore analyzed the transcriptomic profiles of phlebotomized *Erfe-/-* mice 1 and 2 days after phlebotomy compared to control mice.

**Figure 2:**
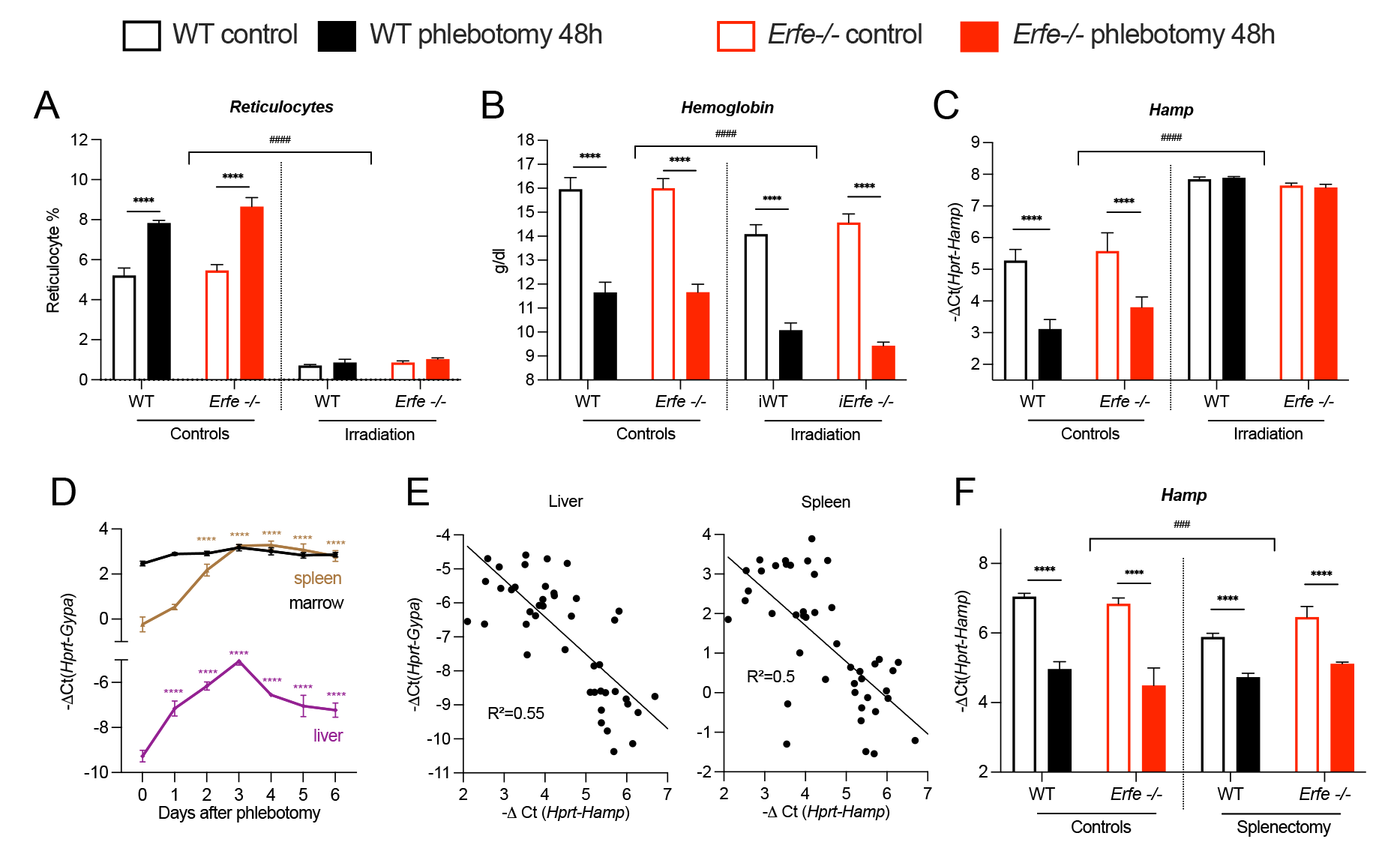
Testing the potential contribution of an erythroid regulator. Percentage of reticulocytes (A), hemoglobin levels (B) and liver *Hamp* (C) mRNA expression in 8 week old control and irradiated (400 rads) WT and *Erfe-/-* mice at t=0 and 48 hours after phlebotomy (n=4-9). (D) *Gypa* mRNA expression in the bone marrow, spleen and liver of *Erfe-/-* at t=0 to 6 days after phlebotomy (n=5-8). (E) Linear regression analysis of *Gypa* and *Hamp* mRNA expression in the liver and the spleen. (F) Liver *Hamp* mRNA expression in 7-9 week-old WT control and splenectomized WT and *Erfe-/-* at t=0 and 48 hours after phlebotomy (n=3-6). Data shown are means ± s.e.m and were compared between each group by Two-way ANOVA. \*\*\*\**P* < 0.0001, \*\*\**P* < 0.001, ^####^*P* < 0.0001 (controls vs irradiated).

### *Fgl1* mRNA expression is induced in mouse livers during anemia

To identify potential regulators of hepcidin, we searched for transcripts encoding secreted proteins and whose expression was induced 24 and 48 hours after phlebotomy compared to control mice. We found and confirmed by qRT-PCR that Fibrinogen-like 1 (FGL1) mRNA expression was significantly induced in the liver and the bone marrow (Figure 3A-B) 1-3 days after phlebotomy in WT and *Erfe-/-* mice but several order of magnitude higher in the liver. FGL1, also known as hepassocin^29^ or HFREP-1^30^, is a member of the fibrinogen family of proteins mostly produced by hepatocytes that share structural homologies to angiopoietin-like proteins (ANGPTL)^31^, including a C-terminal globular domain homologous to fibrinogen beta and gamma subunits. In contrast with other fibrinogen-related factors, FGL1 lacks the platelet-binding and thrombin-sensitive sites involved in clot formation^30, 32^. Instead, FGL1 was induced during liver regeneration and showed mitogenic activity on hepatocytes^33^. It is also involved in tumor evasion of certain cancers through its interaction with LAG-3 receptor^34^. Intra-peritoneal injection of EPO (200u) in WT mice led to a significant reduction in *Hamp* mRNA expression but did not stimulate *Fgl1* expression (Figure 3C). However, *Fgl1* mRNA expression was upregulated in the liver of thalassemic *Th3/+* and *Th3/+* mice deficient for *Erfe* (Figure 3D). To examine whether *Fgl1* expression was stimulated by the decreased oxygen saturation in the liver of anemic mice, we compared *Fgl1* expression in mouse primary hepatocytes incubated in low oxygen conditions (2% O2) or in presence prolyl-hydroxylases inhibitor DMOG to mouse primary hepatocytes incubated under control conditions. We observed that expression of hypoxia inducible factor target genes *Vegfa*, *Gapdh*, *Angptl1* was increased in cells cultured in serum free or serum-containing media and incubated for 15 hours in presence of DMOG or in low oxygen condition (2%) (Supplemental figure 5) compared to untreated cells. Similarly, *Fgl1* mRNA expression was induced in mouse primary hepatocytes incubated in hypoxic conditions or with DMOG (Figure 3E). A trend toward an increase was observed in livers of *Vhlh^lox/lox^/AlbuminCre*^20^ (Figure 3F), and a significant increase in *Fgl1* mRNA expression was detected in livers of hepatocyte specific *Hif2α*-overexpressing mice^35^ and of mice treated chronically with prolyl hydroxylase inhibitor vadadustat^19^ (Figure 3G). These results suggest that *Fgl1* expression may be regulated by hypoxia-inducible factors.

**Figure 3:**
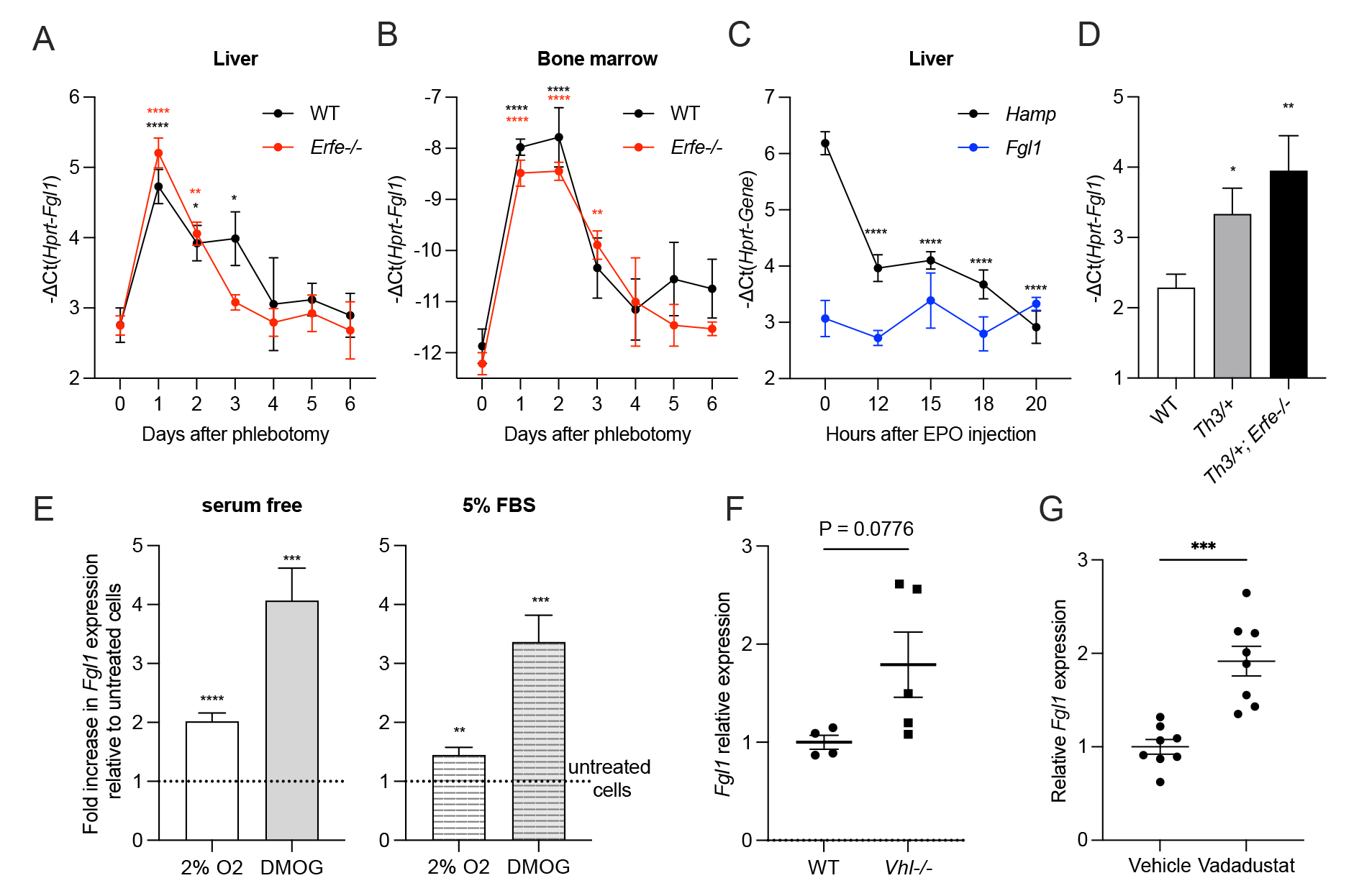
*Fgl1* mRNA expression is induced in mouse liver during anemia. Time course of *Fgl1* mRNA expression in the liver (A) and the bone marrow (B) 1 to 6 days after phlebotomy in WT and *Erfe-/-* mice (n=5-8). (C) *Hamp* and *Fgl1* mRNA expression in the liver of 7 week-old mice at t=0 to 20h after a single intra-peritoneal injection of EPO (200u) (n=5). (D) *Fgl1* mRNA expression in liver of 8 week-old WT, *Th3/+* and *Erfe-/-; Th3/+* mice (n=7-9). (F) Fold change of *Fgl1* mRNA expression in mouse primary hepatocytes in serum free or serum-containing (5% FBS) media and incubated for 15 hours in low oxygen condition (2%) or in presence of prolyl hydroxylases inhibitor DMOG compared to untreated cells (dashed line). Relative *Fgl1* mRNA expression in the liver of *Vhl*-deficient mice (G), and in the liver of mice treated with prolyl hydroxylase inhibitor vadadustat (H). Data shown are means ± s.e.m and were compared for each time point to values for control WT at t = 0 by Two-way ANOVA (A, B, C) or to WT or vehicle treated mice by Student t-test (D, F, G). Data shown for experiment in primary hepatocytes are means of three independent experiments and were compared to control cells by Student t-test (E). \*\*\*\**P* < 0.0001, \*\*\**P* < 0.001, \*\**P* < 0.01, \**P* < 0.05.

### FGL1 is a suppressor of hepcidin *in vivo* and *in vitro*

We next evaluated whether hepatocyte exposure to FGL1 is sufficient for hepcidin suppression. *HAMP* mRNA expression was induced 600 and 200-fold in response to BMP6 (25 ng/ml, 6h) in Hep3B and HepG2 cells respectively (Figure 4A). Treatment with recombinant Fc-tagged FGL1 for 6 hours led to a significant reduction in *HAMP* and *ID1* (Figure 4B-C) expression in presence of BMP6 in both hepatoma cell lines. To confirm these results and identify the active domain of FGL1, we treated Hep3B cells and mouse primary hepatocytes with Fc, full length FGL1 (FL) or its N-terminal (Nter) and globular (glob) domains. We found that full length FGL1 and the globular domain repress *HAMP* and *ID1* expression (Figure 4D-E) whereas the N-terminal domain is inactive. Similarly, injection of recombinant FGL1 (10 mg/kg) into WT mice led to a significant reduction in hepatic *Hamp* RNA expression and serum hepcidin concentration (Figure 4F-G) compared to Fc-treated mice (n=5) 6 hours after injection but no change in liver *Id1* and *Smad7* mRNA expression was observed (Figure 4H-I). Finally, FGL1 was also able to repress hepcidin expression in presence of IL-6 in Hep3B cells (Supplemental figure 6). Collectively, these results indicate that FGL1 is a potent suppressor of hepcidin.

**Figure 4:**
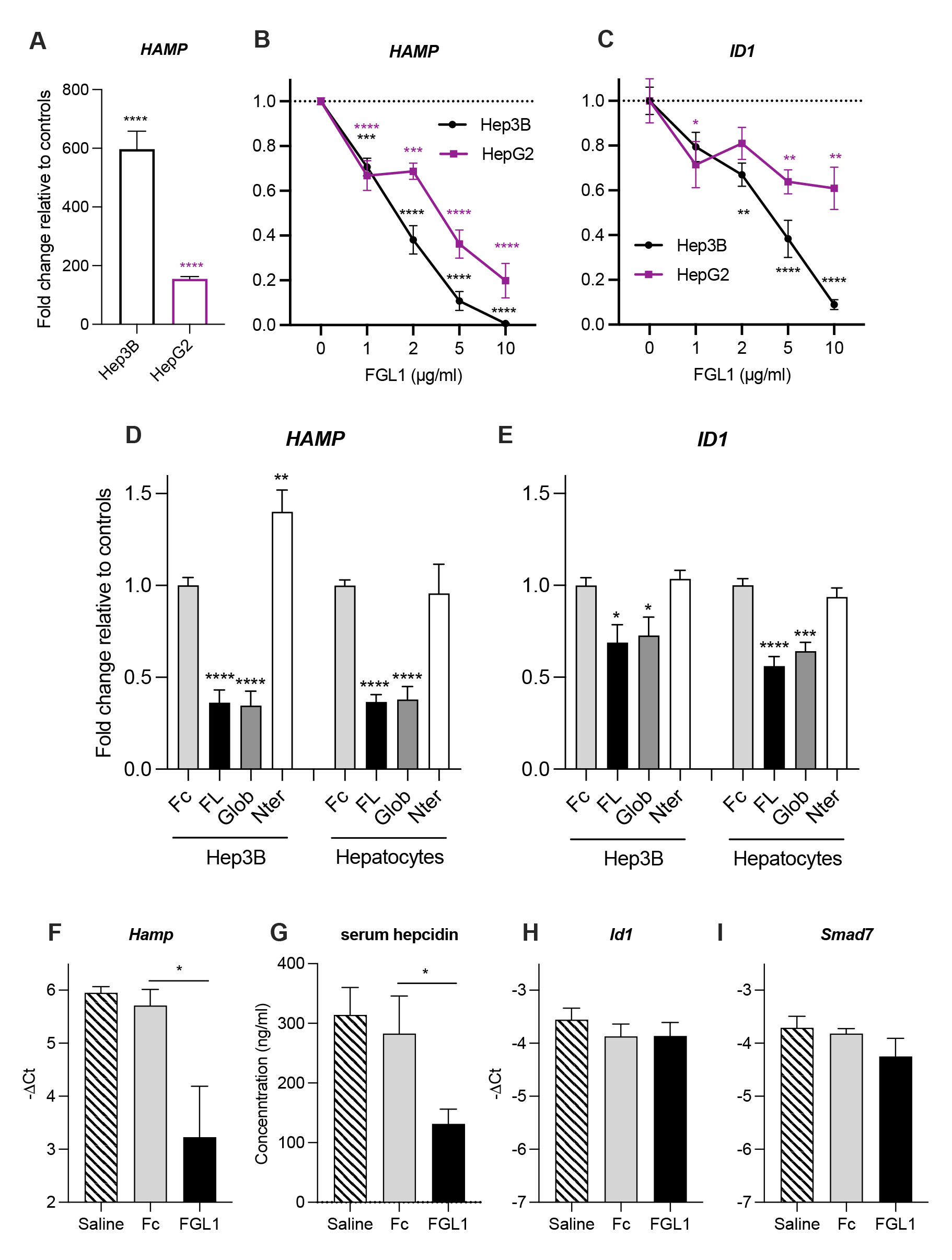
FGL1 is a suppressor of hepcidin *in vivo* and *in vitro*. Relative *HAMP* expression in Hep3B and HepG2 cells in response to BMP6 (25 ng/ml, 6h) (A) or BMP6 + recombinant FGL1 (10µg/ml) (B). Relative *ID1* expression in hepatoma cell lines treated with BMP6 and FGL1 (C). *HAMP* (D) and *ID1* (E) expression in Hep3B cells and mouse primary hepatocytes treated for 6h with BMP6 and either Fc, full length FGL1 and the N-terminal or globular domain of FGL1. Hepatic *Hamp* RNA expression (F), serum hepcidin concentration (G) and liver *Id1* (H) and Smad7 (I) mRNA expression in mice treated for 6 hours with saline, Fc or recombinant FGL1 (10 mg/kg) (n=5). Data shown are means ± s.e.m of three independent experiments (A- E) or treated mice and were compared for each condition to untreated cells or control mice by Student t-test. \*\*\*\**P* < 0.0001, \*\*\**P* < 0.001, \*\**P* < 0.01, \**P* < 0.05.

### *Fgl1-/-* mice exhibit a blunted response to phlebotomy

To determine whether FGL1 contributes to hepcidin regulation during the recovery from anemia, we compared WT and *Fgl1-/-* mice 36 hours after phlebotomy. We first confirmed that *Fgl1* mRNA expression was significantly increased in livers of male and female WT mice 36 hours after bleeding compared to control mice (Figure 5A). Red blood cell count and hemoglobin (Figure 5B-C) levels were reduced 36 hours after phlebotomy in WT and *Fgl1-/-* mice. A comparable increase in *Erfe* mRNA expression in the bone marrow and the spleen was detected 36 hours after bleeding in both genotypes (Figure 5D-E). Interestingly, we found that liver *Hamp* mRNA expression was reduced in male and female phlebotomized WT and *Fgl1-/-* compared to control mice but to a lesser extent in *Fgl1-/-* mice (Figure 5F) suggesting that FGL1 contributes to hepcidin suppression *in vivo*. No significant difference in *Id1* (Figure 5G) and *Smad7* (Figure 5H) mRNA expression was observed between genotypes but only a minor decrease in *Id1* expression was observed in phlebotomized male mice. These results indicate that FGL1 is increased during recovery from anemia and contributes to hepcidin suppression.

**Figure 5:**
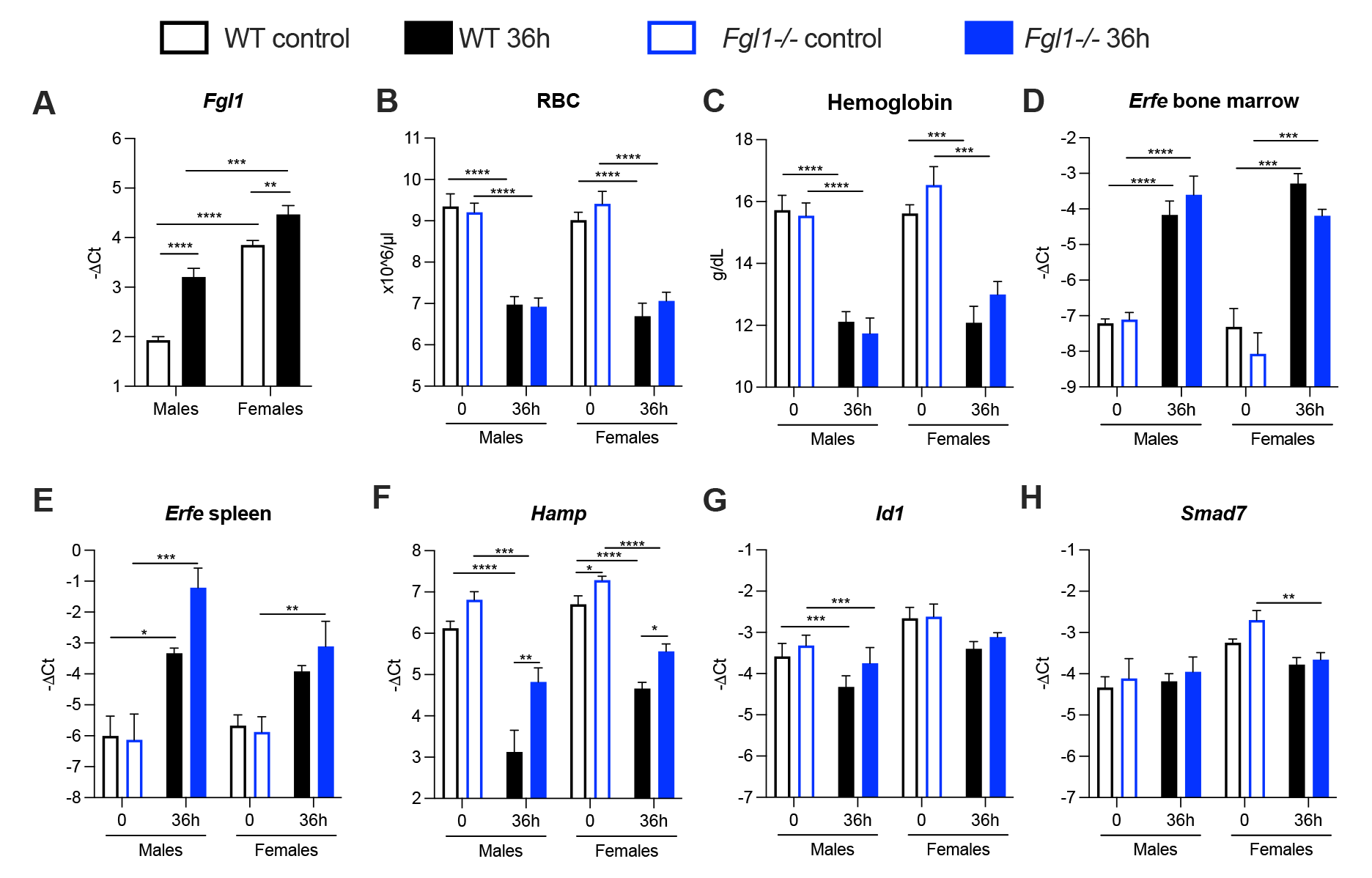
*Fgl1-/-* mice exhibit a blunted response to phlebotomy. *Fgl1* mRNA expression in the liver of male and female WT mice 36 hours after bleeding compared to control mice (A). Red blood cell count (RBC) (B) and hemoglobin (Hb) (C), marrow (D) and spleen (E) *Erfe* mRNA expression, liver *Hamp* (F), *Id1* (G), *Smad7* (H) mRNA expression in WT and *Fgl1-/-* mice at t=0 or 36 hours after phlebotomy. Data shown are means ± s.e.m (n= 5-11) and were compared between each group by Two-way ANOVA and corrected for multiple comparisons by Holm-Šidák method. \*\*\*\**P* < 0.0001, \*\*\**P* < 0.001, \*\**P* < 0.01, \**P* < 0.05.

### FGL1 is a BMP antagonist

We next examined the mechanism by which FGL1 represses hepcidin. Since treatment of hepatic cells with FGL1 leads to a downregulation of *ID1* mRNA expression, we tested whether FGL1 could act as a BMP antagonist. We observed that FGL1 can repress the induction of *Hamp* and *Id1* mRNA expression by BMP 6 and 7 but not by BMP2 and 4 (Figure 6A-B) in mouse primary hepatocytes in comparison to control cells and Fc-treated cells. However, we only observed a small decrease in *Smad7* mRNA expression when cells are treated with BMP7 (Figure 6C). In addition, we found that FGL1 led to a significant reduction in SMAD5 phosphorylation in BMP6-treated Hep3B cells (Figure 6D). Finally, we incubated Fc or Fc-FGL1 (FL, glob and Nter) with BMP6 and performed pull-down assay using protein A magnetic beads. We found that FGL1 FL and glob but not Fc and the N-terminal domain interacted with BMP6 (Figure 6E). Altogether, these results indicate that, similar to erythroferrone, FGL1 may act as a ligand trap for BMP6 to repress hepcidin transcription during the recovery from hemorrhage.

**Figure 6:**
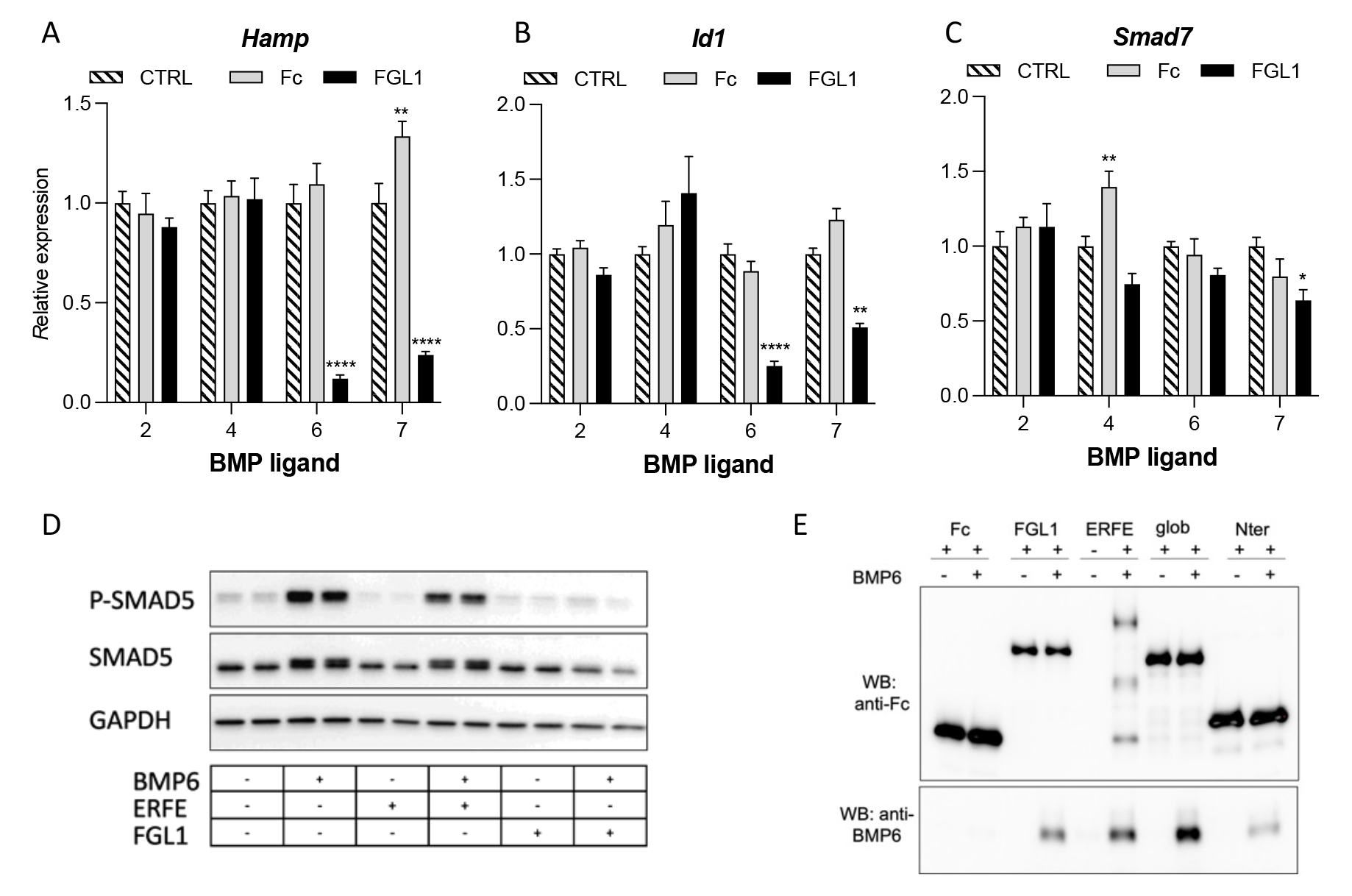
FGL1 is a BMP antagonist. Relative expression of *Hamp* (A), *Id1* (B), *Smad7* (C) mRNA expression in mouse primary hepatocytes treated with BMP ligand (10ng/ml) and human Fc IgG2 (10µg/ml) or Fc-FGL1 (10µg/ml) for 6 hours. Data shown are means ± s.e.m of three independent experiments and were compared for each BMP between Fc or FGL1 treated cells and control cells by Two-way ANOVA. \*\*\*\**P* < 0.0001, \*\*\**P* < 0.001, \*\**P* < 0.01, \**P* < 0.05. (D) Western blotting of Hep3B cells treated for 6 hours with BMP6 (10 ng/ml), ERFE (1µg/ml), or FGL1 (10µg/ml) for P-SMAD5, SMAD5 and GAPDH. (E) Western blotting of pull down assay of BMP6 and FGL1 (FL, glob, Nter) for human Fc IgG2 and BMP6.

## DISCUSSION

The observation that, in humans, iron absorption is regulated by the extent of the iron stores and the rate of erythropoiesis has first been suggested in 1958^36^. When erythropoiesis in the bone marrow intensifies in response to hypoxia-driven synthesis of EPO, iron consumption by erythroid precursors and hemoglobin synthesis can increase up to 5-10 fold^37^. To fulfill the iron needs for erythropoiesis, liver hepcidin synthesis is suppressed by a so called erythron-related regulator^3^ in order to stabilize ferroportin and increase intestinal iron absorption and iron mobilization from stores. Over the last decade, the erythroid hormone erythroferrone (ERFE)^11^ has emerged as the long sought physiological suppressor of hepcidin. Under the influence of EPO, ERFE is secreted by erythroblasts and binds to soluble BMP ligands in the liver perisinusoidal space to prevent the activation of their cognate receptors. This in turn decreases the phosphorylation of SMAD effectors and hepcidin transcription. As a stress hormone, ERFE promotes the recovery from anemia induced by hemorrhage^11^, chronic inflammation^10^, chronic kidney disease^38^ and malarial infection^39^. However, mice deficient for *Erfe* do recover from anemia with a few days delay compared to their WT counterparts. One possible explanation would be that, without the rapid (24h) compensatory suppression of hepcidin by ERFE, increased erythropoietic rate and iron consumption by erythroid precursors result in a transient hypoferremia which, in turn, decreases the activation of the BMP/SMAD signaling and hepcidin transcription^40, 41^. A comparable observation has been made in *Erfe-/-* mice subjected to acute^40^ or chronic EPO treatment^42^. Another likely possibility is that prolonged anemia and hypoxia stimulate the production of a yet-unidentified repressor of hepcidin.

In pathologies associated with ineffective erythropoiesis such as non-transfusion dependent β-thalassemia^12, 16^, myelodysplastic syndromes with ring sideroblasts^13^ or congenital dyserythropoietic anemia^14^, pathological overproduction of ERFE leads to iron overload and severe clinical complications associated with increased mortalitity and morbidity. Indeed, in thalassemic mice, the genetic ablation of ERFE^16^ or its inhibition using neutralizing antibodies^17^ raised hepcidin to levels comparable to those of control WT mice. However, the increase in hepcidin relative to the liver iron content was still blunted compared to WT mice. Indeed, inhibition of ERFE was not sufficient to improve ineffective erythropoiesis and only slightly decreased iron overload further supporting the possibility that an additional hepcidin repressor plays a role during anemia.

Here, we studied the complete recovery from a single blood withdrawal of a third of the total blood volume (500µl) in mice. In our model, the nadir anemia was observed three days after phlebotomy and the mice had recovered after five to six days (Figure 1). Interestingly, hepcidin expression was repressed over 5 days while serum ERFE levels had become undetectable after three days. Our data indicate that an ERFE-independent mechanism contributes to hepcidin regulation between 24-48h after phlebotomy in male and female mice deficient for *Erfe* (Figure 1 and supplemental figure 1). The downregulation of hepcidin was not preceded by any change in liver and serum iron concentration, transferrin saturation, hepatic BMP/SMAD signaling as shown by the phosphorylation status of effectors SMAD1/5/8 (Supplemental figure 3) and the mRNA expression of BMP target genes *Id1* and *Smad7*. Even though hepcidin suppression differed between WT and *Erfe-/-* mice, we did not observe any difference in hemoglobin recovery between WT and *Erfe-/-* mice in contrast with our previous study^11^. In the current experiment, we analyzed a different group of mice at each time point whereas the same mice were previously monitored daily over 9 days^11^. Other parameters such as the genetic background, standard rodent chow, dietary iron content and bioavailabilty, animal facility and housing conditions could influence these discrepancies.

Disruption of the erythroid compartment prevented hepcidin suppression 48 hours after phlebotomy in both WT and *Erfe-/-* mice suggesting that a second erythroid regulator may exist. Competing signals could also hinder hepcidin regulation by erythropoiesis in this model. Indeed, bone marrow ablation leads to decreased reticulocytes and erythroblasts iron consumption which increases serum iron concentration and transferrin saturation^43^ and stimulates the BMP/Smad signaling (Supplemental figure 7-8) and hepcidin expression through the HFE/TFR2 axis^44, 45^. Importantly, the iron signal seems dominant over the erythropoietic drive as ERFE is less efficient in repressing hepcidin in iron-loaded WT mice^11, 42, 46^. Moreover, increased liver *Saa1* mRNA expression indicates that the radiation caused an inflammatory response that could also stimulate hepcidin expression through the IL-6/STAT3 or BMP/SMAD signaling^47–49^.

Extramedullary erythropoiesis is a common feature of patients with chronic anemia (e.g. hereditary spherocytosis, autoimmune hemolytic anemia…) or ineffective erythropoiesis (e.g. thalassemias, myelodysplastic syndromes). Glycophorin A mRNA measurements suggested that extramedullary erythropoiesis in the liver and the spleen was inversely correlated with hepcidin expression 1-3 days after phlebotomy. Of note, hepcidin was still repressed after phlebotomy in splenectomized animals (Figure 2) thus excluding the contribution of a spleen-derived factor.

We therefore initiated a search by microarray for potential hepcidin suppressors derived from the liver and the marrow. We focused specifically on transcripts encoding secreted proteins that were induced 24 and 48 hours after phlebotomy in *Erfe*-deficient mice. We found that Fibrinogen-like 1 (FGL1) mRNA expression was induced in the liver and the marrow 24-48 hours after phlebotomy in both WT and *Erfe-/-* mice. FGL1 is a member of the Fibrinogen superfamily produced mostly by hepatocytes. It binds to fibrin but does not participate in the clotting process^32^. Interestingly, as radiation induces the apoptosis of erythroid precursors and progenitors causing stress erythropoiesis and anemia^50^, liver *Fgl1* expression was already maximally induced 5 days after irradiation (Supplemental figure 8). Its potential contribution in hepcidin regulation after phlebotomy could therefore not be appreciated. This is also consistent with its proposed contribution during hepatic injury, liver failure and liver regeneration^51–53^ as radiation injury is often accompanied by oxidative stress and acute inflammatory responses in irradiated regions^54^. FGL1 has been described as an acute phase protein that is induced by IL-6^51^ but we did not observe a significant increase in IL-6 target gene *Socs3* expression in our phlebotomy model (data not shown). Similarly, our data indicate that *Fgl1* is not an EPO-responsive gene but its expression was increased in the liver of thalassemic mice compared to control mice (Figure 3). We hypothesized that hypoxia could be driving *Fgl1* transcription. In line with this hypothesis, *Fgl1* mRNA expression was increased in the liver of *Hif2α*-overexpressing mice^35^ and in hepatocytes incubated in hypoxic conditions or treated with the prolyl-hydroxylase inhibitor DMOG. A trend toward an increase and a mild yet significant induction of *Fgl1* expression was observed in the liver of *Vhlh^lox/lox^/AlbuminCre* mice and mice treated with the prolyl-hydroxylase inhibitor vadadustat, respectively (Figure 3). However, hypoxia is accompanied by both HIF stabilization and chromatin modifications^55^, a prerequisite to expose the hypoxia-responsive element. Our murine models were restricted to HIF stabilization but chromatin opening may be also required to study the regulation of *Fgl1* by hypoxia.

Treatment of human hepatoma cells lines or primary mouse hepatocytes with recombinant FGL1 resulted in a robust suppression of hepcidin and *Id1* mRNA expression (Figure 4). In contrast, while mice treated with FGL1 exhibited decreased liver and serum hepcidin levels, no significant change in *Id1* and *Smad7* mRNA expression was observed. Interestingly, a crosstalk between fibrinogen and the BMP/SMAD signaling has been reported in oligodendrocyte progenitor cells^56^ and neural stem/precursor cell^57^. In contrast with the BMP agonist property of fibrinogen, we show FGL1 acts as a BMP antagonist that directly binds BMP6 to repress hepcidin transcription (Figure 6). Both ERFE and FGL1 may thus regulate hepcidin through a similar mechanism. However, our data suggest that FGL1 cannot repress the expression of hepcidin induced by BMP2 and BMP4. Interestingly, we did not observe any change in SMAD signaling in phebotomized mice (Figure 1 and S3) and in mice treated with FGL1 (Figure 5). This is in line with previous studies^11, 42, 46, 58^ describing that EPO/ERFE-mediated suppression of hepcidin was not paralleled by a downregulation in *Id1* mRNA expression in mice treated with EPO and in *Erfe*-overexpressing transgenic mice. *In vivo*, the BMP signaling is rapidly feedback-regulated by serum iron concentration and the EPO-induced redistribution of iron. The FGL1-mediated reduction in SMAD signaling may not be discernible in the context of erythropoietic drive, serum and liver concentration, all converging on BMP/SMAD signaling to regulate hepcidin.

*Fgl1*-deficient mice do not harbour any obvious phenotype with the exception of an increased body weight and a glucose intolerance^31^. We did not observe any difference in hematological parameters between WT and *Fgl1-/-* mice at 8 weeks of age (Supplemental Figure 9). Nevertheless, ablation of *Fgl1* may influence baseline hepcidin expression as we observed a mild yet significant increase in hepcidin mRNA expression in female *Fgl1-/-* mice compared to WT mice and a non-signifcant trend toward an increase in male mice (Figure 5). Since the repression of hepcidin occurs between 24 and 48 hours after phlebotomy in *Erfe-/-* mice, we compared the hepcidin response to bleeding in WT and *Fgl1-/-* 36 hours after phlebotomy. While *Erfe* mRNA expression was similarly upregulated in the marrow and spleen of both genotypes, hepcidin suppression was blunted in *Fgl1-/-* mice confirming that FGL1 contributes to hepcidin regulation during the recovery from anemia.

Although the primary goal of this study was to decipher hepcidin regulation between 24 and 48h post-bleeding, we noted that hepcidin expression remained repressed for over 5 days in WT mice, despite both *Erfe* and *Fgl1* mRNA levels returning to baseline 3 to 4 days after phlebotomy. In contrast with ERFE, we could not examine FGL1 protein levels in the liver or the circulation. We tested a large number of commercially available antibodies that all proved unspecific when tested on *Fgl1-/-* samples. Whether the downregulation of hepcidin when hemoglobin levels are nearly recovered after 4 days is mediated by another yet unidentified regulator or by massively increased erythropoiesis and susbsequent iron consumption by erythroblasts remains to be determined.

As a new BMP antagonist that can repress hepcidin even in presence of inflammatory cytokines, FGL1 may be a promising therapeutic candidate to treat anemia of inflammation. Although no validated assays to measure plasma FGL1 have been reported, clinical studies including measurements of plasma FGL1 in patients with chronic anemias will be necessary to determine its contribution in human pathologies.

## Acknowledgements

The author thank members of the INSERM US006 facility (Toulouse) and Janvier labs (Le Genest St Isle) for their technical assistance and help in the mouse breeding, Anthony Emile and Yannick Lippi for their contribution to microarray fingerprints acquisition and microarray data analysis carried out at GeT-TRiX Genopole Toulouse Midi-Pyrénées facility and the platform Aninfimip, an EquipEx (‘Equipement d’Excellence’) supported by the French government through the Investments for the Future programme. Support for this work was provided by the French National Research Agency (ANR-16-ACHN- 0002-01 and ANR-22-CE14-0076-01) and by the European Research Council (ERC) under the European Union’s Horizon 2020 research and innovation program (grant agreement no. 715491) to LK, The French Society for Hematology (SFH) to US, the National Institutes of Health, National Institute of Diabetes and Digestive and Kidney Diseases to T.G. (R01DK126680), the “Fondation pour la Recherche Médicale” EQ202103012630, the Laboratory of Excellence GR-Ex, (ANR-11-LABX-0051), the program “Investissements d’avenir” of the French National Research Agency, reference ANR-11-IDEX-0005-02 to CP, the National Institute of Diabetes and Digestive and Kidney Diseases (NIDDK) (R01 DK107670 and DK095112 to Y.Z.G.).

## Authorship Contributions

US and PP performed experiments and analyzed data, KC, MS, AD, LC, MRM, JT, BB, GJ, EA, performed experiments, CP and YG collected data, EN and TG provided reagents, collected data and edited the manuscript, LK designed and supervised the study, analyzed data and wrote the manuscript.

