## Supplemental data for "The hepatokine FGL1 regulates hepcidin and iron metabolism during the recovery from hemorrhage-induced anemia in mice"

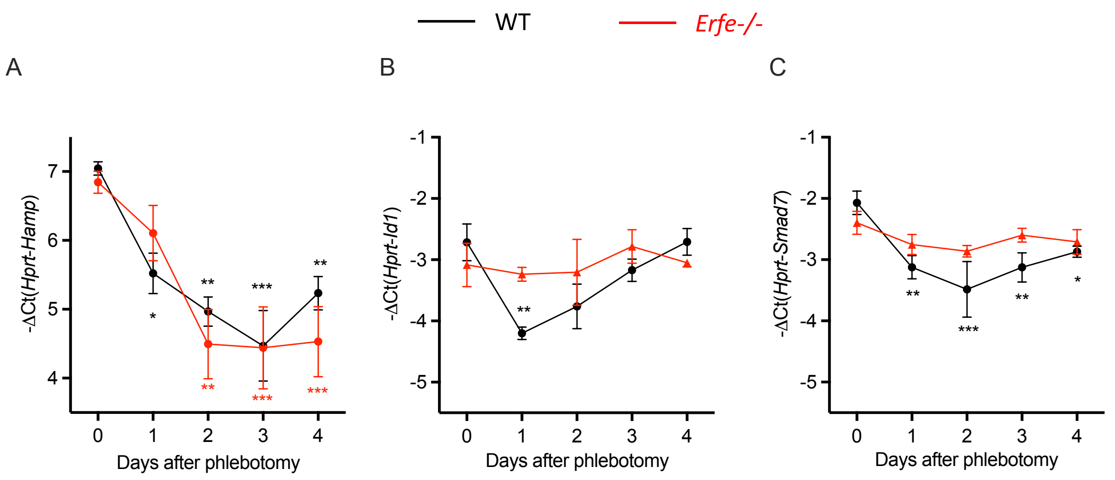


**Supplemental figure 1: ERFE-independent repression of hepcidin during the recovery from anemia.** *Hamp* (A), *Id1* (B) and *Smad7* (C) mRNA expression in the liver of 7-9 week-old WT and *Erfe-/-* female mice 0-4 days after phlebotomy (500µl). Data shown are means ± s.e.m and were compared for each time point to values for control mice at t = 0 (n = 4) by Two-way ANOVA. ***$P < 0.001$, **$P < 0.01$, *$P < 0.05$.

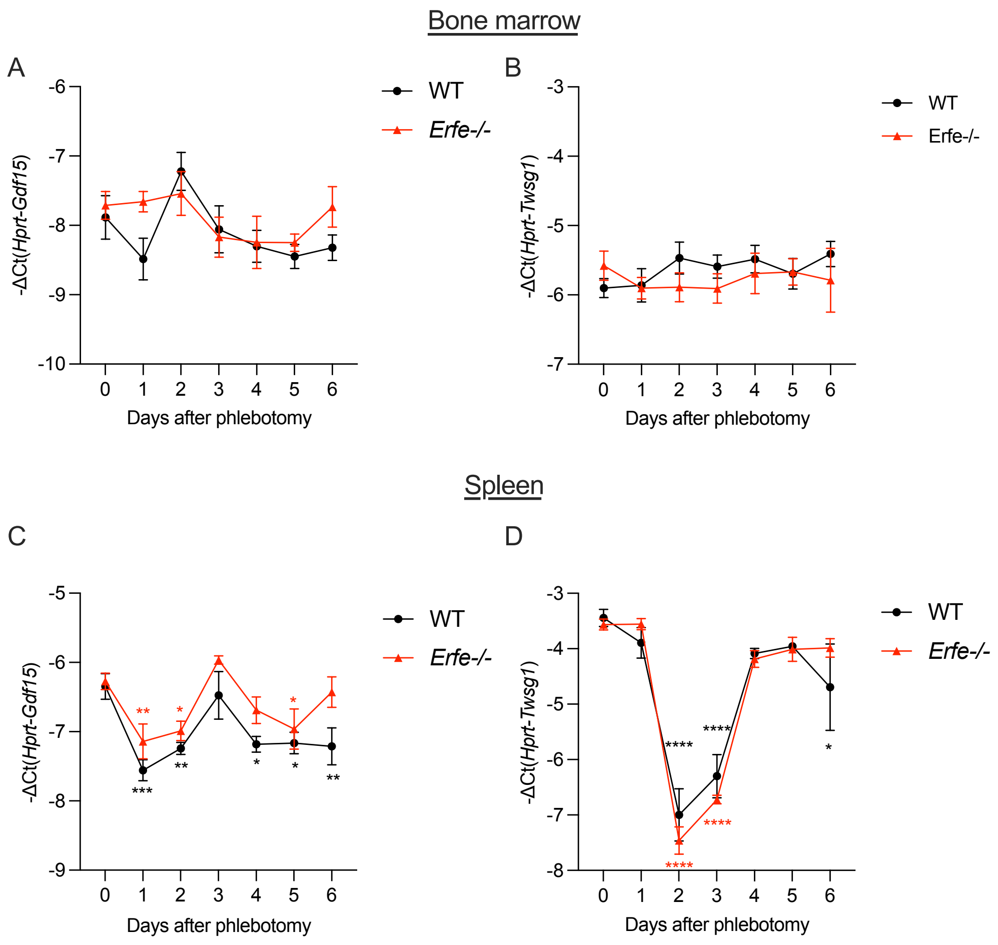


**Supplemental figure 2: GDF15 and TWSG1 are not responsible for hepcidin suppression during the recovery from anemia.** *Gdf15* and *Twsg1* mRNA expression in the liver of 7-9 week-old WT and *Erfe-/-* mice 0-6 days after phlebotomy (500µl). Data shown are means ± s.e.m and were compared for each time point to values for control mice at t = 0 (n = 5-8) by Two-way ANOVA. ****$P < 0.001$, ***$P < 0.001$, **$P < 0.01$, *$P < 0.05$.

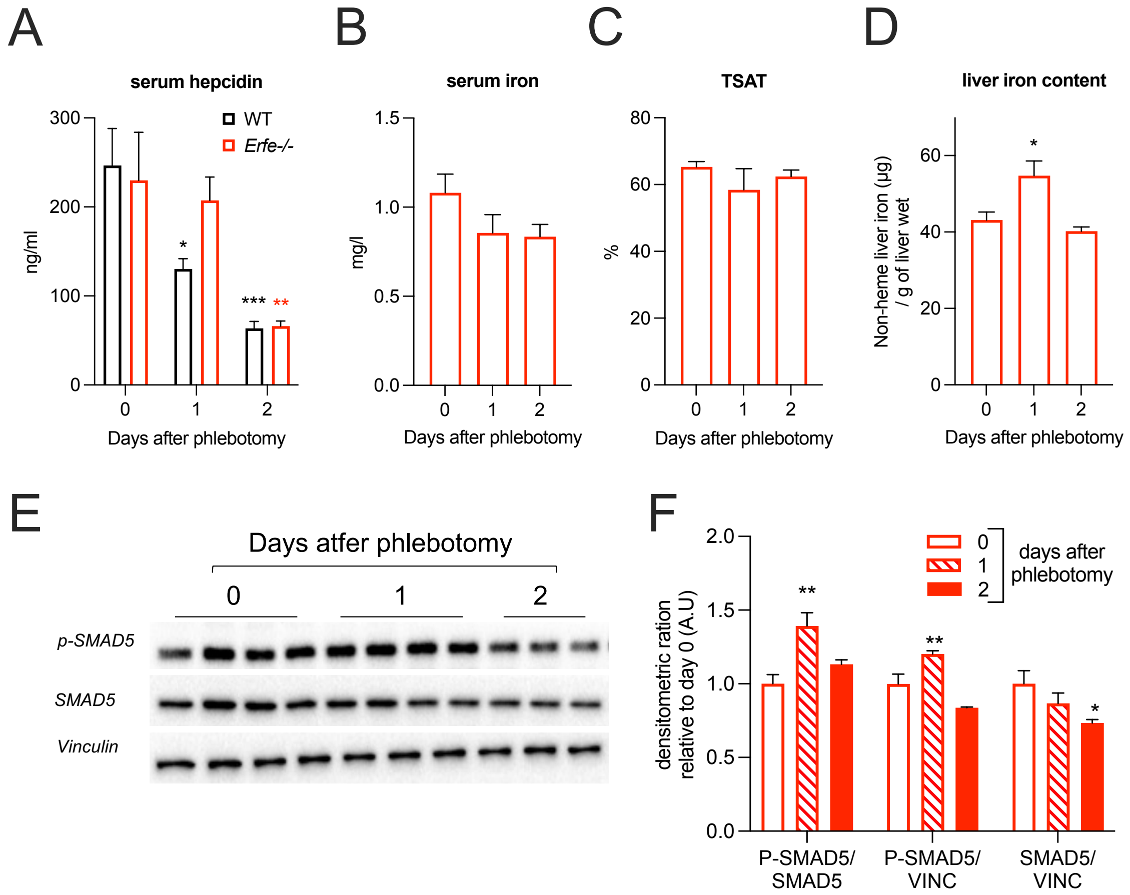


**Supplemental figure 3: ERFE-independent repression of hepcidin during the recovery from anemia is not mediated by iron parameters.** Serum hepcidin concentration (A), serum iron content (B), transferrin saturation (C) and liver iron content (D) in 7-9 week-old *Erfe-/-* mice 0-2 days after phlebotomy (500µl). (E) Western blotting for P-Smad5, Smad5 and vinculin in the liver of *Erfe-/-* mice 0, 1 and 2 days after phlebotomy. (F) Densitometric ratio of phosphorylated Smad5 to total Smad5 or Vinculin and Smad5 to Vinculin. Data shown are means ± s.e.m and were compared for each time point to values for control mice at t = 0 (n = 5-8) by Student t-test. ****P* < 0.001, ***P* < 0.01, **P* < 0.05.

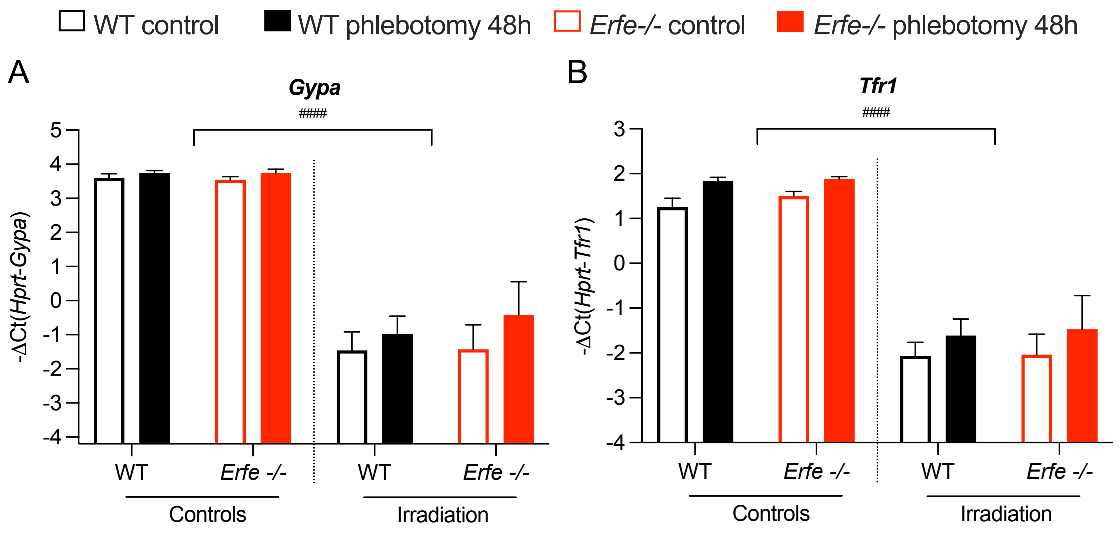


**Supplemental figure 4: Glycophorin A and transferrin receptor 1 mRNA expression in the marrow of control and irradiated mice.** *Gypa* (A) and *Tfr1* (B) mRNA expression in the bone marrow of 8 week-old control and irradiated (400 rads) WT and *Erfe-/-* mice at t=0 and 48 hours after phlebotomy (n=4-9). Data shown are means ± s.e.m and were compared between each group by Two-way ANOVA. $^{\#\#\#\#}P < 0.0001$ (controls vs irradiated).

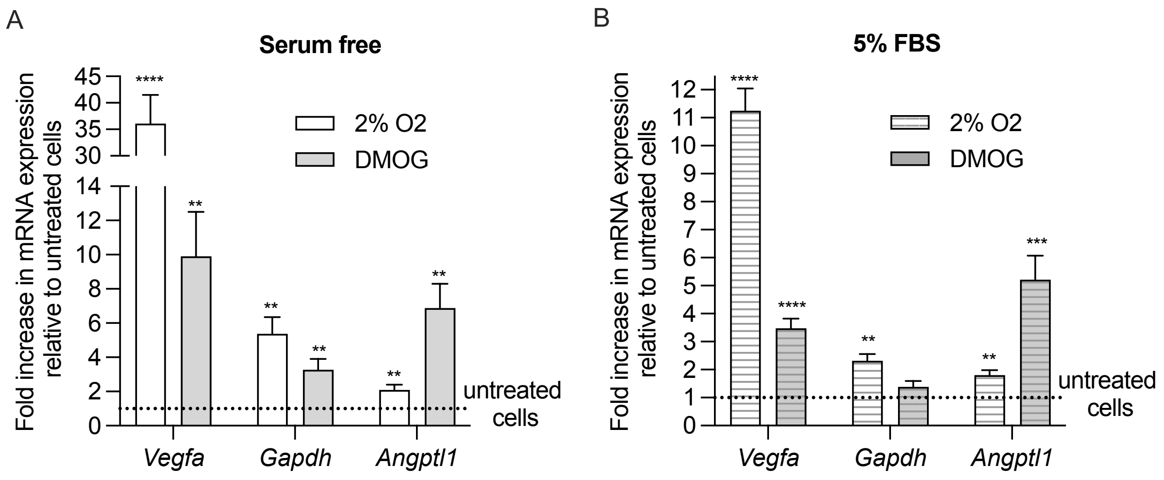


**Supplemental figure 5: Effect of hypoxia (2% O2) and prolyl hydroxylases inhibitors DMOG on hypoxia responsive genes in mouse primary hepatocytes.** Fold change in mRNA expression of HIF target genes *Vegfa*, *Gapdh* and *Angptl1* in mouse primary hepatocytes cultured in serum free (A) or serum-containing (B) media and incubated for 15 hours in low oxygen condition (2%) or in presence of prolyl hydroxylases inhibitor DMOG compared to untreated cells (dashed line). Data shown are means of three independent experiments and were compared to untreated control cells by Student t-test. $****P < 0.0001$, $***P < 0.001$, $**P < 0.01$, $*P < 0.05$.

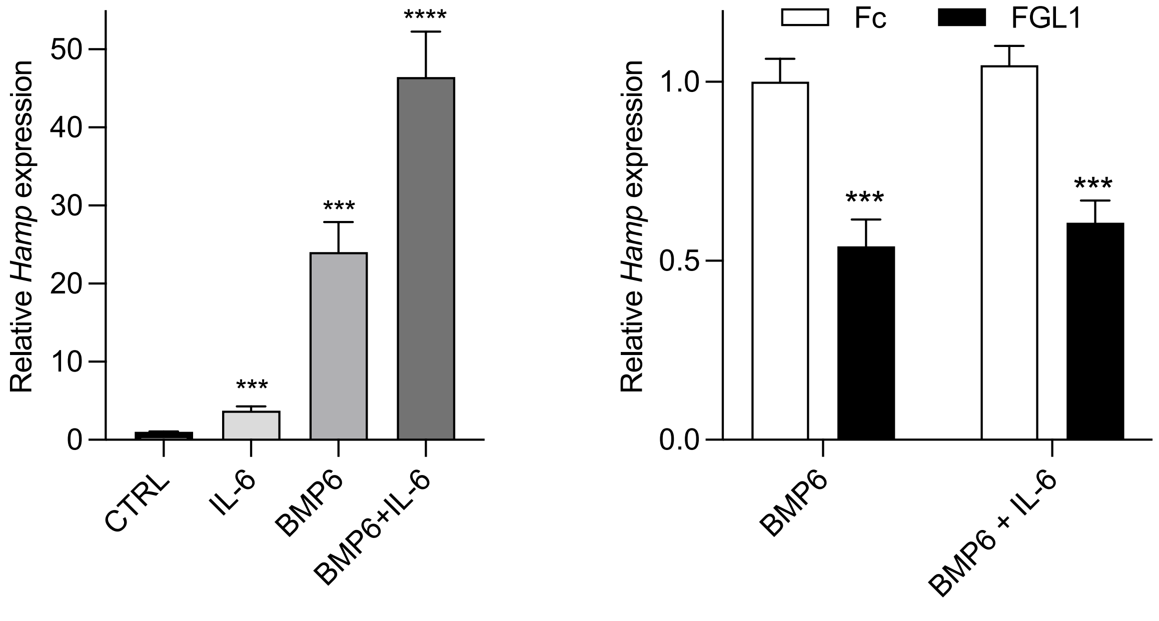


**Supplemental figure 6: FGL1 can repress hepcidin in presence of IL-6.** Relative *Hamp* expression in mouse primary hepatocytes in response to BMP6 (25 ng/ml, 6h), IL-6 (20 ng/ml) or BMP6+IL-6 (A) and in presence of Fc and FGL1 (10µg/ml) (B). Data shown are means ± s.e.m of three independent experiments and were compared for each condition to untreated (A) or Fc-treated (B) cells. ****$P < 0.0001$, ***$P < 0.001$.

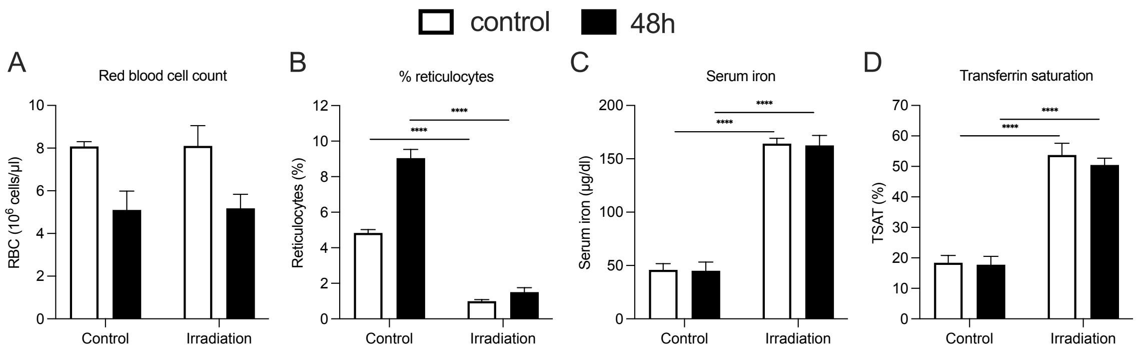


**Supplemental figure 7: The effect of irradiation on blood and iron parameters.** Red blood cell count (A), reticulocytes percentage (B), serum iron concentration (C) and transferrin saturation (D) in 7 week-old control and irradiated C57Bl/6J mice at t=0 and 48 hours after phlebotomy (n=5). Data shown are means ± s.e.m and were compared between each group by Two-way ANOVA and were corrected for multiple comparisons by Holm-Šidák method. ****$P < 0.0001$.

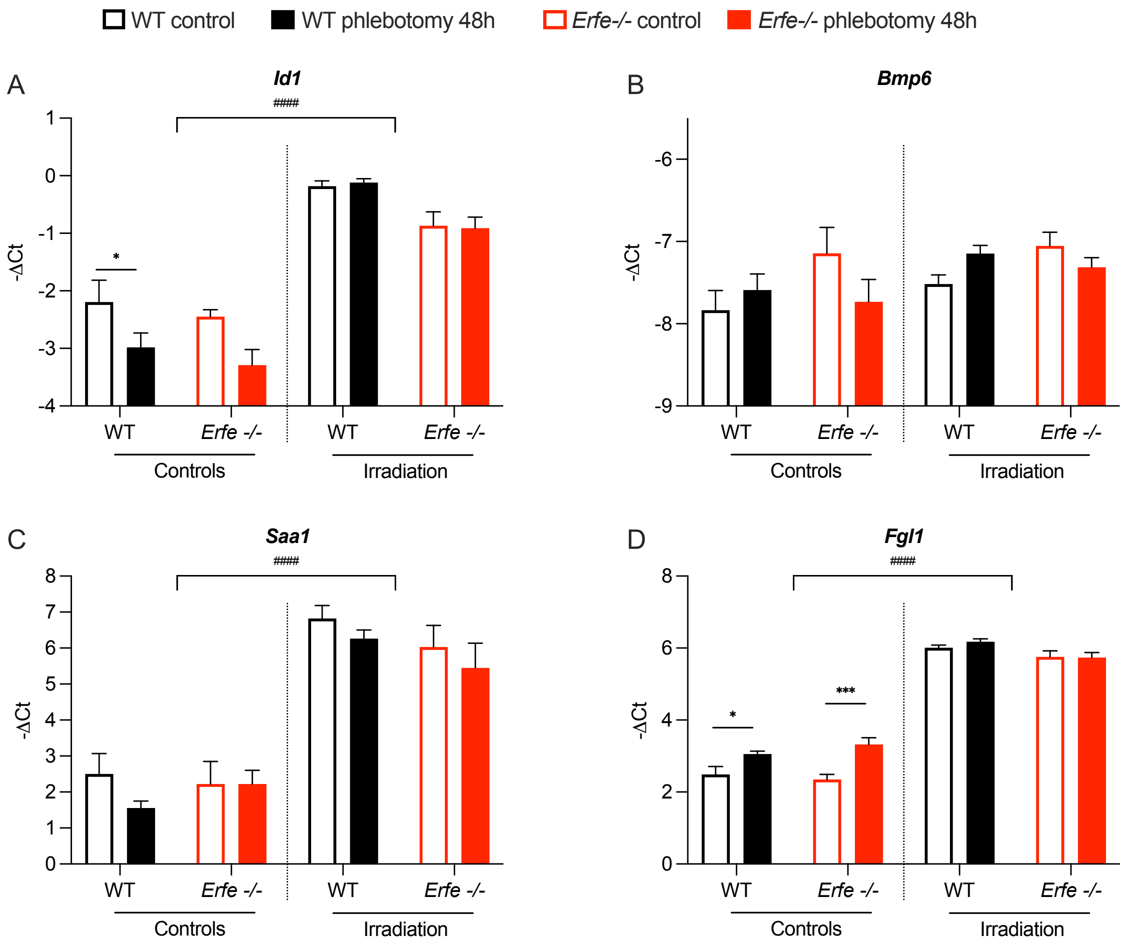


**Supplemental figure 8: The effect of irradiation on the liver.** Liver *Id1* (A), *Bmp6* (B), *Saa1* (C) and *Fgl1* (D) mRNA expression in 8 week-old control and irradiated (400 radian) WT and *Erfe-/-* mice at t=0 and 48 hours after phlebotomy (n=4-9). Data shown are means ± s.e.m and were compared between each group by Two-way ANOVA and were corrected for multiple comparisons by Holm-Šidák method. ****P* < 0.001, **P* < 0.05, $^{\#\#\#\#}$*P* < 0.0001 (controls vs irradiated).

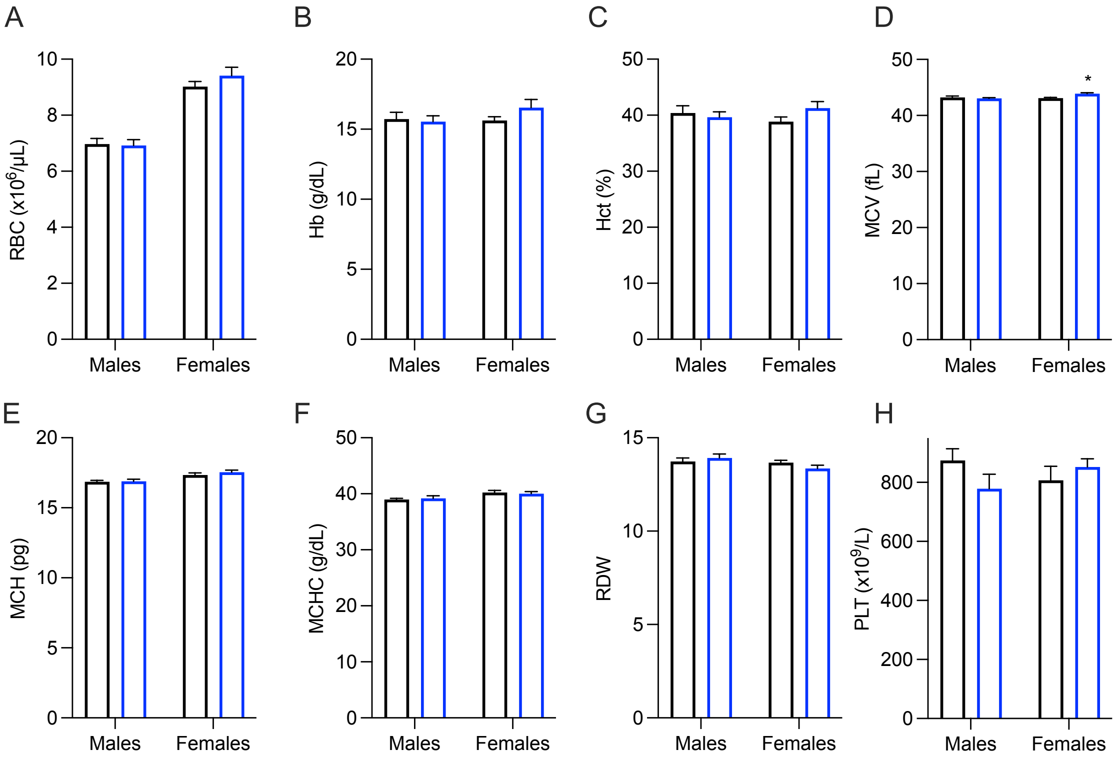


**Supplemental figure 9: Complete blood count in WT and *Fgl1-/-* mice.** Red blood cell count (A), hemoglobin (B), hematocrit (C), mean corpuscular volume (D), mean corpuscular hemoglobin (E), mean corpuscular hemoglobin concentration (F), reticulocyte distribution width(G) and platelet count (H) in 8 week-old WT and *Fgl1-/-* mice (n=8-11). Data shown are means ± s.e.m and were compared between WT and *Fgl1-/-* mice by Two-way ANOVA and corrected for multiple comparisons by Holm-Šidák method. **P* < 0.05.

**Supplemental table 1:**

| Gene name | Primer Forward | Primer Reverse |
|---|---|---|
| *hHPRT* | AAGCTTGCGACCTTGACCAT | TGCTTTCCTTGGTCAGGCAG |
| *hHAMP* | CCAGCTGGATGCCCATGTT | GCCGCAGCAGAAAATGCA |
| *hID1* | CCCTCAACGGCGAGATCAG | GTTTCCCCGTCGGTATAAGGA |
| *hFGL1* | GATCAGTCTGGCTGGTGGTT | TGCCAGGTGTACCAGACAAT |
| *mHprt* | CTGGTTAAGCAGTACAGCCCCAA | CGAGAGGTCCTTTTCACCAGC |
| *mHamp* | GCAGGGCAGACATTGCGAT | AGAGAGGTCAGGATGTGGCTCT |
| *mId1* | ACCCTGAACGGCGAGATCA | TCGTCGGCTGGAACACATG |

| | | |
|---|---|---|
| *mSmad 7* | GCAGGCTGTCCAGATGCTGT | GATCCCCAGGCTCCAGAAGA |
| *mAtoh8* | CAGAAGGGCGAGCCAAGAAACG G | CTGGTGGTCCCAGCTTTCTCCTC A |
| *mBmp6* | ATGGCAGGACTGGATCATTGC | CCATCACAGTAGTTGGCAGCGT |
| *mGdf15* | GCTGTCCGGATACTCAGTCCA | TTGACGCGGAGTAGCAGCT |
| *mTwsg1* | AGCGACAAAGAGCGCATGTG | CACTGGTGGATGGACATGCAG |
| *mGypA* | GGAGGAATGCCGTCACCAA | TAATCCCTGCCATCACGCC |
| *mTfr1* | CACGAGCGGAATACAGCCA | CCCATGACGTTGAATTGAACCT |
| *mEpo* | GCCTCACTTCACTGCTTCGG | GGAGGCGACATCAATTCCTTC |
| *mErfe* | ATGGGGCTGGAGAACAGC | TGGCATTGTCCAAGAAGACA |
| *mFgl1* | CGATCTGATGGCAGTGAGAACT | TTTGTTACCCAGCCAGTATTCG |
